# PanXpress: Gene expression quantification with a pan-transcriptomic gapped k-mer index

**DOI:** 10.64898/2026.03.19.712873

**Authors:** Inês Alves Ferreira, Jens Zentgraf, Johanna Elena Schmitz, Sven Rahmann

## Abstract

**Motivation:** Most existing workflows for quantifying bacterial gene expression from RNA-seq data rely on mapping reads to a (single) reference transcriptome, typically ignoring strain-level variation. When samples contain unknown or mixed strains, these workflows may introduce reference bias and fail to accurately capture strain-specific gene expression. Pan-transcriptomic approaches address this issue by using pan-transcriptomes as references, but existing solutions require multiple steps for pan-transcriptome construction, indexing, and expression quantification.

**Results:** We introduce PanXpress, a unified framework for bacterial pantranscriptomics that performs pantranscriptome construction and indexing directly from genomic FASTA and GFF annotation files, alignment-free mapping of reads to genes from FASTQ samples, and gene expression quantification. The index, a multi-way Cuckoo hash table storing gapped *k*-mers with associated genes, preserves diversity on the *k*-mer level.

Using simulated RNA-seq data from a mixture of Pseudomonas *aeruginosa* strains, PanXpress achieves mapping recall comparable to alignment-based methods such as Bowtie2 with higher precision and obtains accurate gene expression and log fold change estimates. On real *P. aeruginosa* RNA-seq data, using PanXpress’ pantranscriptomic reference increases the proportion of mapped reads and discovered expressed genes. The index of PanXpress is smaller than that of other tools and it provides faster analysis with consistent results, compared to other tools (Salmon, Kallisto, Bowtie2). PanXpress is thus an accurate and efficient method for bacterial gene expression analysis in complex samples.

**Availability:** PanXpress is available at https://gitlab.com/rahmannlab/panxpress.

## 1 Introduction

Accurate quantification of bacterial gene expression is essential for understanding antibiotic resistance [14]. Standard RNA-seq analysis workflows rely on a single reference genome, typically from a dominant or well-studied strain. However, reads from unknown or mixed strains may map poorly or remain unmapped, making it necessary to incorporate genomic variation across strains. Pan-transcriptomic approaches address this by using a pan-transcriptome as the reference.

Two challenges arise in pan-transcriptomic analysis. First, we must represent variation within strain-specific sequences. Second, we must group orthologous genes across strains, complicated by paralogs arising from gene duplications. Existing tools address the second challenge via construction of classical pangenomes [9], dividing genes into core genes (present in all strains) and accessory genes (present in a subset of strains), with additional classifications, such as soft core (≥95% of genomes) [11] or rare genes (≤15%) [20]. Pangenome construction is typically a preprocessing step, performed using tools such as Panaroo [19], PPanGGOLiN [6], Roary [13], or PIRATE [1]. They output a consensus sequence per gene, losing strain-specific nucleotide variation, and require separate read mapping using alignment-based methods, such as Bowtie2 [8], or pseudo-alignment (alignment-free) methods, such as Salmon [16] or Kallisto [2], followed by expression quantification.

The above approach has several disadvantages. The mentioned pangenome tools are designed primarily for comparative genomics and population-level analyses. They need specific annotation formats and generate extensive reports describing gene families, presence–absence matrices, and classifications of genes as core or accessory. While valuable for pangenome analysis studies, this information is not required for transcriptomic read mapping, and the associated algorithms introduce substantial computational overhead. The selection of a single representative sequence per gene cluster discards strain-specific nucleotide variation, rendering the output less useful for transcriptomic read mapping. In summary, no existing framework inte grates pan-transcriptome construction, indexing, and gene expression inference in one method.

We introduce PanXpress, a new approach that integrates pan-transcriptome construction, read-to-gene mapping, and gene expression quantification directly from user-provided genome FASTA sequence files and GFF annotation files. It enables rapid mapping by constructing a gapped *k*-mer index over the pantranscriptome reference. Gene expression levels are estimated by counting reads mapped to each gene and normalizing counts to transcripts per million (TPM). For differential expression analysis between different conditions, PanXpress converts read-mapping outputs into count matrices compatible with PyDESeq2. An overview of PanXpress is shown in Figure 1.

**Figure 1.**
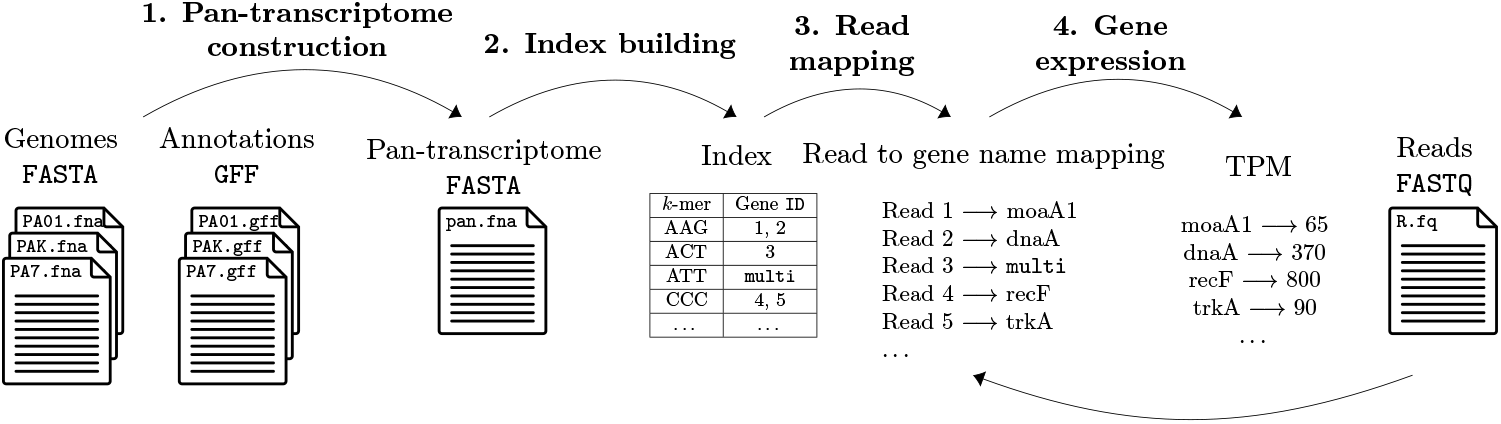
Overview of PanXpress. **1. Pan-transcriptome construction**: To construct the pan-transcriptome reference (output), the user inputs the genome `fasta` files of the bacterial strains and the respective annotation files. **2. Index building**: We now use the pan-transcriptome reference as an input to build an index where each *k*-mer in the reference is matched to the gene(s) where it is present. **3. Read mapping**: With the index and the reads `fastq` files, we can find a gene match for each read. **4. Gene expression**: With the read mapping information, we can count how many reads are matched to each gene and output the gene expression information.

This work is organized as follows. Section 2 introduces necessary definitions. Section 3 explains pantranscriptome construction (Sec. 3.1), index creation (Sec. 3.2), and read mapping (Sec. 3.3). Section 4 presents the experimental setup and experimental results, including comparisons to Bowtie2, Salmon, and Kallisto. Section 5 provides concluding remarks.

## 2 Preliminaries

We introduce basic concepts related to *k*-mers. A DNA sequence *s* is a string over the DNA alphabet Σ = {*A, C, G, T*} . The length of *s* is written as |*s*| . Let rc(*s*) denote the reverse complement of *s*, obtained by reversing *s* and replacing each nucleotide by its complement. A *k*-mer of *s* is a substring of length *k* of *s*. A string of length *n* contains (*n− k* + 1) *k*-mers, some of which may be equal. To efficiently represent *k*-mers, we encode each nucleotide using a 2-bit representation, so a *k*-mer can be represented as a 2*k*-bit integer. Given a *k*-mer, we compute both its encoding and the encoding of its reverse complement. We define the canonical representation of both the *k*-mer and its reverse complement as the *maximum* of these two integer values. This ensures that a *k*-mer and its reverse complement are treated identically.

A *gapped k-mer* is a string of length *k* obtained by selecting *k* specific positions within a window of length *w≥ k*. The selected positions, also called *significant positions*, are specified by a *mask*. A (*k, w*)-mask is a string of length *w* over the alphabet {`#`, `_`} that contains `#` exactly *k* times and `_` exactly (*w − k*) times. The positions marked by `#` are the significant ones and concatenated to yield a gapped *k*-mer, whereas positions marked by `_` are ignored and are referred to as “don’t care positions”. Thus, a gapped *k*-mer of *s* is a specific subsequence of any length-*w* substring of *s*.

It has been shown in the past that gapped *k*mers can be more robust to genomic variation, in particular to single nucleotide variants (SNVs), than standard contiguous *k*-mers [22]. For example, if 4 substitutions are arbitrarily distributed on a sequence of length 100, all contiguous 25-mers can be changed (by distributing the changes approximately equidistantly), but using the (25, 35) mask `####### ### #_###_# ### #######` guarantees that at least 4 gapped 25-mers are unchanged, independently of the distribution of the substitutions [22].

The similarity between two sets *A* and *B* can be measured by their Jaccard index 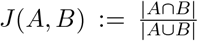. We use the Jaccard similarity to estimate the similarity between two protein sequences based on their amino acid *k*-mer content. This approach does not require an alignment of the protein sequences, and *J* (*A, B*) can be efficiently computed, but *J* (*A, B*) = 1 does not mean that the protein sequences are identical, only that their *k*-mer content agrees.

## 3 Methods

We describe the procedure used by PanXpress for constructing the pan-transcriptome, aiming to retain all nucleotide sequence variants for each gene across strains to allow for accurate read mapping. The main challenges when constructing a pan-transcriptome across strains are dealing with inconsistent gene and protein annotations, especially concerning paralogous genes, and a large number of unnamed “hypothetical protein”s. The first step of PanXpress consists of resolving annotation inconsistencies and paralog ambiguity as much as possible (Sec. 3.1), partitioning annotated genes across strains into groups, with the idea that one group represents all variants of the same gene. Then, all nucleotide sequences of the a group are collected under a common numerical identifier with an associated gene name, yielding a pan-transcriptome nucleotide FASTA file (point 1. in Figure 1). The implementation supports bacterial genomes with multiple chromosomes and plasmids, as well as genes spanning multiple CDS features. Each (gapped) *k*-mer of these sequences is then indexed and associated to the group(s) in which it occurs using a fast hash table (Sec. 3.2). Read-to-gene mapping is done by examining the read’s gapped *k*-mer occurrences across genes and taking a majority consensus vote (Sec. 3.3).

### 3.1 Annotation harmonization

Paralogous genes (“paralogs”) arise from duplication events and may retain, lose, or diverge in function. Unfortunately, their genomic annotation is sometimes inconsistent for the purpose of collecting all variants of the same gene under a common identifier. For example, in the *Pseudomonas aeruginosa* strain PAK (see Supplement for accession numbers), two genes annotated as `moaA` (`WP_003083204.1`, `WP_003092949.1`) share only 45.4% sequence identity, showing they are distant paralogs. In strain PAO1, the same proteins are annotated as `moaA1` (`NP_252559.1`) and `moaA2` (`NP_250196.1`), each matching one of the PAK sequences. Inconsistent naming leads to *k*-mers mapping to multiple identifiers, causing multi-mapping (see Section 3.2). Another inconsistency occurs when different strains assign different gene names to the same protein. In *P. aeruginosa* strain UCBPP-PA14, protein `WP_003090351.1` is annotated as uvrY, whereas in strain LESB58, the same protein is named gacA, with 99.5% sequence identity. Without correction, reads map ambiguously. We proceed in three steps.

#### Step 1

Proteins are conceptually represented as nodes in an undirected graph. Edges connect nodes that represent the same gene, so connected components define protein or gene groups. In order to quickly merge connected components, we use a union-find data structure on the nodes and apply the union-find algorithm [4]. Initially, each annotated coding gene (or protein) in the GFF annotation files forms its own group. This means that identical copies of genes that occur in several strains are present as multiple nodes. Genes occurring on plasmids are renamed by appending `Plasmid` to their names in order to distinguish chromosomal and plasmid copies of the same gene, which can be important for functional analysis [3]. In the first step, nodes that share exactly the same annotated gene name or protein ID are connected.

The following steps aim to connect nodes where the names may differ but the sequence similarity gives sufficient indication that they represent variants of the same gene. Whenever nodes with different gene names are connected, the group name becomes the concatenation of the contained gene names (e.g., `moaA_moaA1`), ignoring “hypothetical protein” to avoid non-descriptive long names.

#### Step 2

We compute the Jaccard similarity of all pairs of proteins, including hypothetical proteins, in all strains based on amino acid 7-mers to discover candidates that we might want to group together based on sequence similarity. Protein-level similarity is preferred over nucleotide-level because proteins capture functional similarity and reduce computational cost, as nucleotide sequences are three times longer and have larger *k*-mer sets. Users may provide protein FASTA files or generate them from GFF annotation and genomic FASTA using tools like AGAT [5]. To reduce memory and runtime, we convert amino acid sequences to a 15-letter reduced alphabet: (L,V,I,M), C, A, G, S, T, P, (F,Y), W, E, D, N, Q, (K,R), H. This preserves biochemical similarity [17]. We chose *k* = 7 to construct *k*-mer sets over this alphabet to balance specificity and sensitivity; *k* = 7 has been reported to capture strain-level similarity reliably [18].

Each protein sequence is converted into a sorted numerical representation of its constituent reduced-alphabet 7-mers. As log_2_(15^7^) *≈*27.3*≤* 32, each 7-mer can be represented by a 32-bit integer. The number of common 7-mers between two proteins is then quickly computed by parallel scanning the two sorted sequences, similar to a merge-sort step.

#### Step 3

Protein pairs with Jaccard similarity exceeding a threshold *t*_1_ are candidates for further examination and scored using standard overlap alignment (using BLOSUM62 scores, a gap open penalty of *−*11 and a gap extension penalty of *−*1; the BLAST defaults),which allows for partial overlaps or containment relations between the sequences. This is relevant for proteins of different lengths, including plasmids and truncated or hypothetical proteins. We compute a normalized score between 0 and 1 by dividing the resulting alignment score by the minimum of their respective self-scores, where a self-score is the score obtained when aligning a protein with itself. Pairs with normalized scores above a threshold *t*_2_ are assigned to the same gene family. Singleton hypothetical proteins are labeled `unnamed`.

#### Rationale for the three-step procedure

Step 1 assumes that there are no annotation errors in the sense that if gene names are identical, the sequences represent the same gene. Step 2 ensures that proteins are only grouped together if they share a certain global similarity on the *k*-mer level. This avoids that short fragments (that match a substring of sequences of several other groups) act as bridges and erroneously merge unrelated groups. Step 3 ensures that sequence similarity with respect to the shorter sequence is sufficiently high. The combination of steps 2 and 3 with appropriate selection of thresholds *t*_1_ and *t*_2_ avoids spurious merges. Threshold selection is detailed in the Supplement.

### 3.2 Gapped k-mer index

PanXpress uses a three-way bucketed Cuckoo hash table [24] that maps each gapped *k*-mer in the re-annotated pan-transcriptome to a set of gene group identifiers, referred to as its *color set*. This variant of Cuckoo hashing allows both relatively fast construction at high load factors (few empty slots) and fast lookups. Figure 2 explains the main idea behind this Cuckoo hashing variant.

**Figure 2.**
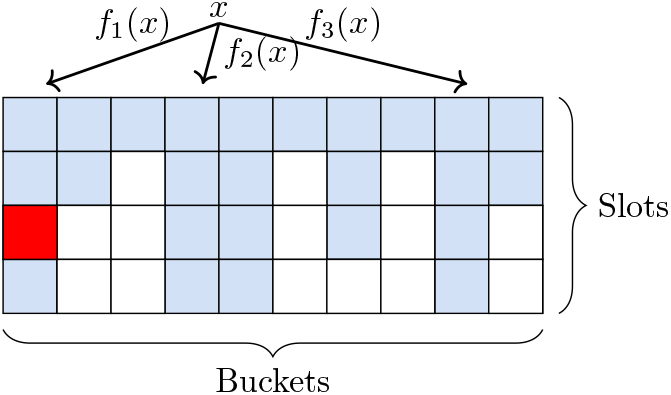
Three-way Cuckoo hash table. Three hash functions, *f*_1_(*x*), *f*_2_(*x*), and *f*_3_(*x*) map an element *x* to three candidate buckets, each containing four slots, yielding 3*×* 4 = 12 possible insertion positions. Blue cells indicate occupied slots. If no slot is available for *x*, one of the 12 positions is chosen uniformly at random (red), the occupying element is evicted, and *x* is inserted. Then re-insertion of the evicted element is attempted at one of its alternative locations. This may be iterated, initiating a random-walk insertion process that continues until success or a predefined failure limit is reached.

The size of the color set is the same for all *k*-mers; the limit is flexible in the implementation but needs to be specified when building the index. We use 4 colors in the following. This choice provides a good tradeoff between genome coverage and memory usage (see Results). Using a small size limit saves us from the complexities of a general flexible color set implementation. If a gapped *k*-mer occurs in more than 4 genes, a special `multi` value is stored instead, indicating that the color set exceeds the allowed capacity.

After all gapped *k*-mers have been inserted into the hash table, they are classified according to their uniqueness properties. A gapped *k*-mer is *unique* if its stored color set contains only one gene identifier; otherwise, it is *non-unique*. Unique gapped *k*-mers are further subdivided into *strongly unique* and *weakly unique* based on their robustness to sequencing errors or single-nucleotide variants. Specifically, a unique gapped *k*-mer is *weakly unique* if there exists at least one Hamming-distance-1 neighbor whose color set differs from its own. In such cases, a single nucleotide substitution may alter the gene assignment, making the *k*-mer less reliable for gene mapping. To capture this property, we store an additional bit per *k*-mer, referred to as the *weak bit*, which indicates whether such a neighboring *k*-mer exists. If no Hamming-distance-1 neighbor with a different color set is found, the unique *k*-mer is classified as *strongly unique*. All pairs of stored gapped *k*-mers at Hamming-distance-1 are efficiently identified using the `Fourway` algorithm [23], which efficiently computes all weak bits without exhaustively comparing all pairs of *k*-mers.

The used Cuckoo hash table has a theoretical load threshold of 99%, but it is faster at a load of 95%. In order to achieve a precise load factor, the number of *k*-mers that will be inserted needs to be known exactly or reliably estimated. We build the table twice; first with a large size and small load to get the exact number; then again with the exact target load factor. For the initial pass, we use the total number of *k*-mers in the FASTA file as an upper bound, Σ_*i*_ (*𝓁*_*i*_*−k* + 1), where *𝓁*_*i*_ is the length of the *i*-th nucleotide sequence in the file. Since *k*-mers are often repeated in variants of the same gene, the actual number is typically much smaller.

### 3.3 Read mapping

Read-to-gene mapping is performed as follows. For each read, we iterate over its gapped *k*-mers. Each *k*-mer is queried in the index to retrieve the associated gene identifiers (if any). All retrieved gene identifiers are collected in a list and counted according to the following criteria: If a *k*-mer is strongly unique, the identifier is counted 5 times; if it is weakly unique, the identifier is counted 3 times; if it is non-unique, each of the up to 4 identifiers is counted once. Absent or `multi` *k*-mers are not counted. After processing all *k*-mers, the collected identifiers are aggregated and sorted by frequency. Let *y*_1_, *y*_2_ be the named gene identifiers, ignoring the unnamed hypothetical proteins, with the highest and second highest frequencies *f*_1_*≥f*_2_. (If *y*_2_ does not exist, we use *f*_2_ = 0 and leave *y*_2_ undefined. Ties where *f*_1_ = *f*_2_ can be resolved arbitrarily without influencing relevant results, as this will result in an unmapped or ambiguously mapped read below.)

If both *f*_1_ *≥T* := 5 absolutely and *f*_1_ *≥f*_2_ + *T* relatively, we say that we have a clear signal for gene *y*_1_ and map the read to it, considering the read successfully *mapped*.

If *f*_1_ *<T*, we check if the frequency of the hypothetical protein group is at least *f*_1_ + *T*, and if this is the case, we assign the read to this class, which does not distinguish between different hypothetical proteins, as no additional information on them is available. Such a read is also considered *mapped*.

If *f*_1_ *< T*, but the frequency of the hypothetical protein class is smaller than *f*_1_ + *T*, the read is considered *unmapped*.

If *T≤ f*_1_ *< f*_2_ + *T*, we argue that we cannot make an unambiguous decision between *y*_1_ and *y*_2_ and consider the read as *ambiguously mapped*.

The above rules apply in this form to single-end reads. We also implemented a paired-end mode, in which the gapped *k*-mers from both reads are combined in the same list and used jointly to determine a mapping decision for the entire read pair, using the same rules as above.

## 4 Results

We compare PanXpress with Kallisto, Salmon, and Bowtie2, using both simulated and real data from *Pseudomonas aeruginosa*. Simulated datasets provide ground truth for benchmarking, allowing evaluation of precision and recall in read-to-gene mapping. For real datasets, true gene expression is unknown, so we compare PanXpress with Kallisto and Salmon in terms of gene expression estimates, and additionally with Bowtie2 for mapping performance, speed and memory usage. We use a publicly available transcriptomic dataset comprising samples from multiple *P. aeruginosa* strains that was previously analyzed to study the relationship between core metabolism and bacterial virulence across strains [15].

Section 4.1 describes our evaluation setup and tool versions. Section 4.2 provides pan-transcriptome and index statistics. In Section 4.3, we present results on simulated reads, and in Section 4.4, we provide results about the real dataset.

### 4.1 Setup and parameters

Benchmarks were run on an AMD Ryzen 9 5950X 16-core CPU (32 threads) with 64 GB RAM, and NVMe data SSDs (WD_BLACK SN850X 4 TB) under Ubuntu 24.04.3 LTS.

For PanXpress, the hash table was constructed with a load factor of 0.95, and we used the following (25, 35) mask described by [22]: `####_###_####_# # #_####_###_####`.

We used Bowtie2 2.5.5 with default parameters. It requires unique FASTA headers, but our pan-transcriptome FASTA reference contains multiple sequences per gene to capture strain diversity, so we created a modified version with unique headers, mapping them back to original identifiers after alignment.

Kallisto v0.51.1 also requires unique headers, which is handled automatically with `–make-unique`. Parameters *l* and *s* (fragment length mean and standard deviation) were set to 100 and 20, respectively. The *k*-mer length was set to 25 to match PanXpress. Salmon v1.10.3 also used the reference with unique headers and back-mapping with 25-mers and parameters `libtype=A`, `fldMean=100`, and `fldSD=20`.

### 4.2 Pan-transcriptome and index

We used the same pan-transcriptomic reference for both simulated and real data. Indices were built separately for each tool. To simulate a realistic scenario, we collected 50 strains of *P. aeruginosa* using the NCBI Datasets CLI tool [12]. The reference strain PAO1 (`GCF_000006765.1`) was included, and 49 strains were randomly selected from complete RefSeq assemblies with annotations (see Supplement for accessions).

We observed that the number of distinct named genes saturates as we add new annotated strains (see Supplement). However, the number of hypothetical proteins increased with each new strain.

The annotation files downloaded from NCBI were harmonized according to the procedure in Section 3.1, using protein FASTA files also downloaded from the NCBI and the thresholds *t*_1_ = 0.02 for Jaccard similarity and *t*_2_ = 0.75 for the normalized alignment score. The choice of these thresholds was informed by a study presented in the Supplement. The above-mentioned dataset of 50 strains was used to build the gapped *k*-mer index that was used for mapping both the simulated and real reads.

We examined the distribution of the number of colors (genes) over indexed *k*-mers to determine a good choice for the maximum color set size (maximum number of gene identifiers that can be stored for each *k*-mer). For any given number of strains (1–50), we randomly sampled that number of strains from the 50 available ones and built the corresponding index, repeating this 10 times with different random sets of strains. The results are shown in Figure 3. Most *k*-mers are unique to a gene or correspond to hypothetical proteins grouped under `unnamed` (*k*-mers shared by hypothetical proteins are counted as having one color). Fewer than 10 *k*-mers are associated with more than four colors, and this barely changes with an increasing number of strains. Therefore, we selected 4 as the upper limit of the color set size to build the PanXpress index. This result is potentially of more general interest: Depending on the application, storing a small color set may suffice most of the time, making a general color set data structure unnecessary, greatly simplifying the implementation. We also see that after including about 40 strains, we only obtain a small number of additional *k*-mers with each new strain.

**Figure 3.**
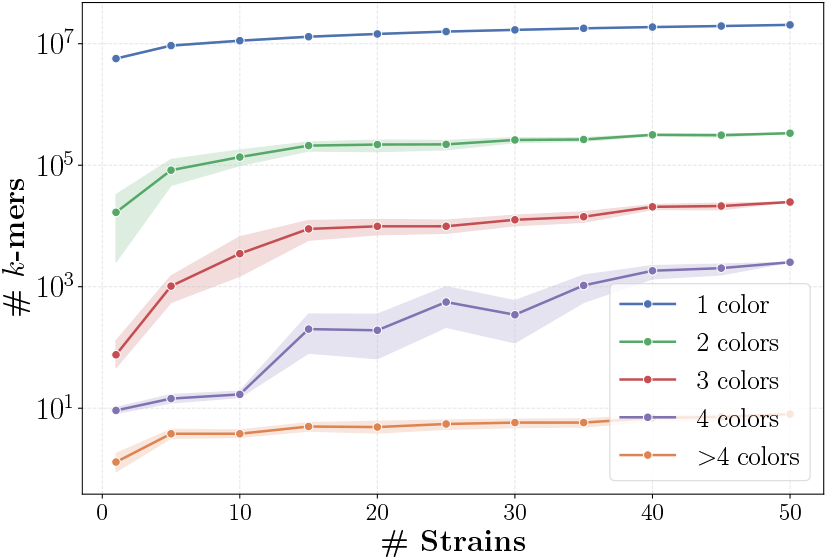
Number of *k*-mers associated with different numbers of colors in the PanXpress index as a function of the number of strains. For each number of strains, the experiment was repeated 10 times using randomly sampled strains. Lines represent the mean number of *k*-mers and the shaded areas indicate 95% confidence intervals.

Concerning index size, PanXpress has the smallest index (206 MB), storing 20.81 million gapped 25-mers and associated gene identifiers in a hash table. Salmon’s Pufferfish index, combining a compacted de Bruijn graph of 25-mers with a minimal perfect hash function, requires 252 MB, while Kallisto’s compacted de Bruijn graph needs 443 MB for the 25-mers with auxiliary structures. Bowtie2’s FM index needs 433 MB of space on the same data.

### 4.3 Simulated data

Simulated reads were generated from a mixture of three strains: PAO1 and two randomly selected ones from the 49 used in the reference. Only named genes were used to simulate reads with ART ILLUMINA [7] using the HiSeq 2500 profile, 100 bp read length, 100*×* coverage, in both single-end and paired-end modes. Two control samples with unmodified expression and two samples with modified expression (15% of genes upregulated ×2; 15% downregulated ×2; remainder unchanged) were generated.

Because read origins are known, we computed mapping precision and recall for PanXpress and Bowtie2 relative to the ground truth (Table 1). PanXpress has (almost) no unmapped reads in this simulated dataset; however, between 0.6% and 0.7% map ambiguously on single-end (SE) data and 0.35% to 0.4% on paired-end (PE) data, explaining recalls slightly below 100%. Notably, almost all of the mapped reads are mapped correctly (100.000% to 3 digits). Bowtie2 leaves almost no reads unmapped or ambiguously mapped, so precision and recall look identical, but about 0.5% (SE) or 0.4% (PE) of the reads are mapped to the wrong gene. Gene-level expression estimates were obtained by counting reads per gene, normalizing by gene length, and analyzing log_2_ fold changes with `PyDESeq2` between the 2+2 control and up-down datasets. Performance of the four methods PanXpress, Bowtie2, Kallisto, and Salmon was compared using root mean squared error (RMSE) and mean absolute error (MAE) between true and estimated fold changes for all named genes (Table 2; detailed plots in the Supplement). Overall, the estimates were accurate, with log_2_ fold change RMSE in the range of 10^*−*3^ and MAE in the range of 10^*−*4^. Salmon achieved the lowest error values for the single-end reads. PanXpress showed the second lowest errors, while the errors of Kallisto and Bowtie2 were higher. On the paired-end dataset, PanXpress performed best, but comparably with Salmon; errors of Kallisto and Bowtie2 again were higher. The improved performance of PanXpress on PE data may be partially explained by the observed better mapping recall on PE data in comparison to SE data (Table 1).

**Table 1.** Read mapping performance comparison between PanXpress and Bowtie2 on simulated *P. aeruginosa* reads, in single-end (SE) and paired-end (PE) mode. Results are reported for two independent control samples (control 1, 2) and two independent samples with modified expression (up-down 1, 2). Precision (Prec%): correctly mapped reads of mapped reads. Recall (Recall%): correctly mapped reads of all reads.

| Dataset | PanXpress |  | Bowtie2 |  |
| --- | --- | --- | --- | --- |
|  | Prec% | Recall% | Prec% | Recall% |
| SE control 1 | 100.000 | 99.330 | 99.510 | 99.510 |
| SE control 2 | 100.000 | 99.334 | 99.512 | 99.512 |
| SE up-down 1 | 100.000 | 99.402 | 99.546 | 99.546 |
| SE up-down 2 | 100.000 | 99.402 | 99.546 | 99.546 |
| PE control 1 | 100.000 | 99.608 | 99.624 | 99.624 |
| PE control 2 | 100.000 | 99.607 | 99.630 | 99.630 |
| PE up-down 1 | 100.000 | 99.650 | 99.652 | 99.652 |
| PE up-down 2 | 100.000 | 99.651 | 99.651 | 99.651 |

**Table 2.** Quantification error comparison for PanXpress, Salmon, Kallisto, and Bowtie2 on simulated *P. aeruginosa* single-end (SE) and paired-end (PE) reads, measured as root mean squared error (RMSE) and mean absolute error (MAE) between estimated and true gene expression log_2_ fold changes.

| Tool | SE RMSE | SE MAE | PE RMSE | PE MAE |
| --- | --- | --- | --- | --- |
| | ($\times 10^{-3}$) | ($\times 10^{-4}$) | ($\times 10^{-3}$) | ($\times 10^{-4}$) |
| PanXpress | 5.528 | 5.026 | **2.763** | **2.393** |
| Salmon | **1.169** | **0.893** | 3.285 | 3.301 |
| Kallisto | 10.909 | 13.845 | 6.178 | 20.829 |
| Bowtie2 | 9.950 | 10.711 | 6.849 | 8.778 |

### 4.4 Real data

We obtained FASTQ reads from the dataset described by [15] (NCBI GEO accession `GSE142464`) containing five *Pseudomonas aeruginosa* strains: PA14, MTB-1, B136-33, CF5, and PACS2. All samples belonging to the same strain were concatenated and treated as a single sample. We additionally obtained the genomic FASTA and GFF annotation files of these strains (accessions in the Supplement). The study by Panayidou et al. [15] used a sequencing protocol that resulted in a high proportion of ribosomal RNA (rRNA) reads. To remove this contamination, we filtered the raw reads using *Cleanifier* [24], a fast *k*-mer-based tool for removing contamination from sequencing data, using an rRNA database containing all sequences available in the SILVA database [21, release 138.2].

We first examined the fraction of mapped reads in each of the five samples obtained when using differently diverse reference transcriptomes (Figure 4 left). For three of the samples (PA14, MTB-1, B136-33), the fraction of mapped reads is lowest when using the standard reference strain PAO1 (A) and only slightly higher when using the single correct reference strain. For the other two samples (CF4, PACS2), this relation is surprisingly reversed, and the fraction of mapped reads is markedly lower when using the correct reference strain, which might be explained by the less complete annotation of that strain compared to PAO1. For all five samples, using the 50-strain pan-transcriptomic reference markedly increases the fraction of mapped reads, and further adding the 5 sample strains to the reference only has a very small additional positive effect. The pan-transcriptomic reference captures a broader set of gene variants, accessory genes, and hypothetical proteins, providing additional mapping targets for reads that would otherwise remain unmapped when using a single-strain reference.

**Figure 4.**
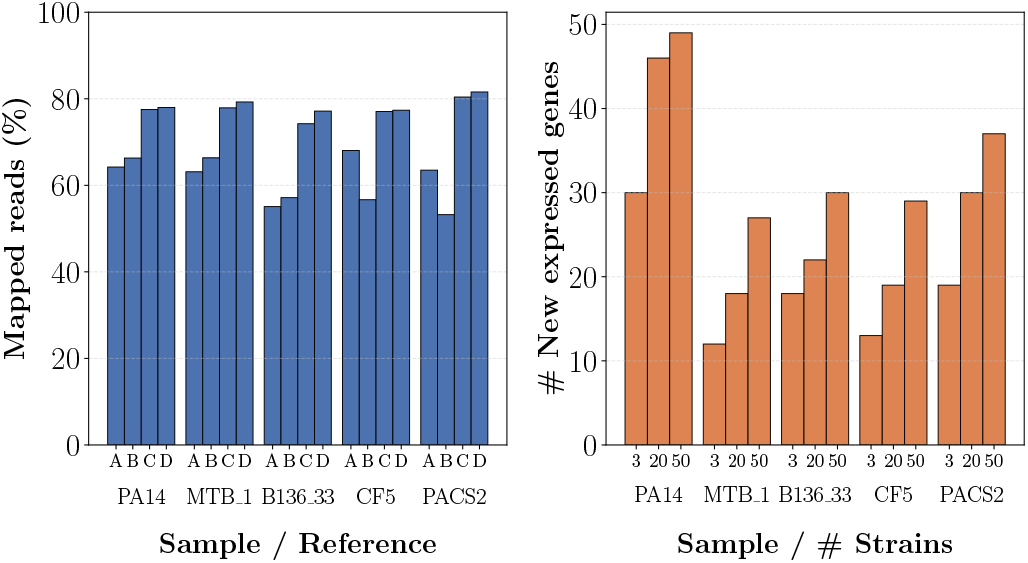
**Left/blue:** Fraction of mapped reads when using different reference transcriptomes ABCD. A: single strain PAO1 (best annotated). B: single true strain from which the reads originate. C: pan-transcriptomic 50-strain reference. D: pan-transcriptomic 55-strain reference (C, plus the 5 strains from the data: PA14, MTB-1, B136-33, CF5, and PACS2). **Right/orange:** Number of additionally detected expressed genes as a function of the number of strains included in the reference pan-transcriptome (3, 20, 50), in comparison to the single-strain PAO1 reference.

We found that a considerable part of the additionally mapped reads map to hypothetical proteins, which might indicate gaps in the annotations. However, we also observed more expressed genes (here defined as genes with a read count of at least 5) in comparison to using the PAO1 strain only (Figure 4 right). As the number of included strains increases (3; 20; all 50 strains), we observe a consistent increase in the number of newly detected expressed genes across all datasets. Although the increase is modest relative to the total of over 2000 expressed genes, identifying these additional genes can be biologically important in the context of antibiotic resistance: One of the genes detected only when using the expanded reference is *istA*, which is absent from and cannot be detected with the standard reference strain POA1. The *istA* gene encodes the transposase of the IS21 family of insertion sequences, which mediate DNA transposition and contribute to genome plasticity. IS21 elements can facilitate the mobilization and spread of antibiotic resistance and virulence genes [10].

Finally, we compared the relative analysis running times between the different tools, fixing the (slowest) time for Bowtie2 with 4 threads to 1.0 for each sample. Relative speed-up factors per sample and tool and for different numbers of threads are shown in Figure 5, measured using wall-clock time (`panxpress map`; `salmon quant`; `kallisto quant`; `bowtie2` mapping step). All tools benefit from more threads, but speedup is slightly sub-linear with increasing thread count. Among the alignment-free methods, PanXpress is fastest by some margin.

**Figure 5.**
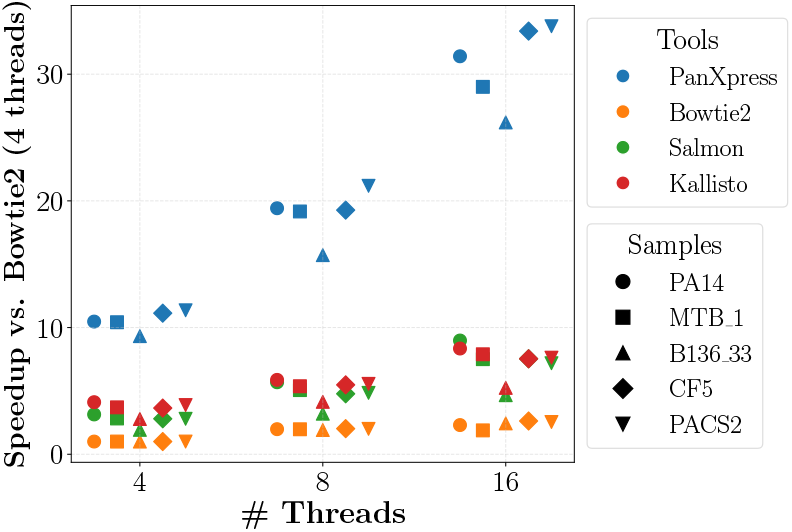
Relative speedup (per sample, tool, and number of threads) in comparison to Bowtie2 with 4 threads (fixed at 1.0 for each sample); higher is faster.

## 5 Discussion and Conclusion

We presented PanXpress, the first unified tool for pan-transcriptome construction, read-to-gene mapping, and gene expression quantification. We developed a novel method for pan-transcriptome construction that addresses common issues with gene annotation across strains in the existing GFF files concerning paralog handling and annotation of unnamed genes that are very similar to named genes in other strains. While our grouping method may lose transcript resolution, we argue that it is more helpful than harmful for its intended purpose, the transcriptomic analysis of samples containing unknown or multiple strains where no reference strain is known or annotated. Expression analysis at the coarse gene level (instead of for specific transcripts) seems appropriate in this situation.

We note that “four colors suffice” for resolving genes in 50 *P. aeruginosa* strains. Extrapolation suggests that this remains true for more strains, but it needs to be investigated systematically for other bacterial species. A study on *Mycobacterium tuberculosis*, presented in the Supplement, shows the same result.

On simulated datasets, we verified that our method can recover planted up- and down-regulation and that read count results are mostly comparable with other alignment-free tools, such as Kallisto or Salmon, as well as alignment-based mappings from Bowtie2. Both index size and speed of PanXpress compare favorably. On real data, using many reference strains instead of a single one yields more mapped reads and more discovered expressed genes, which may be of biological importance.

## A Datasets and Accession Numbers

Table 3 contains the accession numbers for all strains included in the 50-strain *Pseudomonas aeruginosa* dataset, and in addition, for the 5 strains present in the *P. aeruginosa* reads analyzed in the real data section of the results. Table 4 contains the accession numbers for all strains included in the 50-strain *Mycobacterium tuberculosis* dataset. The first entry in each table corresponds to the reference strain of the bacteria.

**Table 3.** NCBI accession numbers and corresponding *Pseudomonas aeruginosa* strain names. The 5 strains at the bottom are the ones used in the analysis of the real dataset.

| NCBI accession | strain name |
| --- | --- |
| GCF_000006765.1 | **PAO1** |
| GCF_001722025.1 | PA_D5 |
| GCF_002753655.1 | 12939 |
| GCF_002812845.1 | PB368 |
| GCF_002968655.1 | AR_0353 |
| GCF_003369735.1 | Y89 |
| GCF_003369775.1 | Y31 |
| GCF_003429205.1 | AES1M |
| GCF_003798105.1 | H26027 |
| GCF_004355125.1 | AES1M |
| GCF_013260445.1 | YD001 |
| GCF_013393665.2 | SE5331 |
| GCF_013394455.1 | SE5443 |
| GCF_016105885.1 | TJ2014-049 |
| GCF_016864415.1 | PARM801 |
| GCF_019857485.1 | ZPPH1 |
| GCF_020990465.1 | P4970C |
| GCF_021497345.2 | SE5381 |
| GCF_021513455.1 | R04 |
| GCF_022570475.1 | H02 |
| GCF_022670795.1 | PA1_NCHU |
| GCF_023101265.1 | AR19640 |
| GCF_024300845.1 | R20-14 |
| GCF_024507955.1 | ATCC 27853 |
| GCF_025723085.1 | Pa3 |
| GCF_027570755.1 | PALA23 |
| GCF_027570815.1 | PALA29 |
| GCF_029961285.1 | 2021CK-01161 |
| GCF_029961305.1 | 2021CK-01158 |
| GCF_029962165.1 | 2022CK-00068 |
| GCF_030034615.1 | ZY94 |
| GCF_030121855.1 | NY5506 |
| GCF_030122015.1 | NY5530 |
| GCF_030122055.1 | NY5535 |
| GCF_030253475.1 | 22112 |
| GCF_033217895.1 | HPA2120 |
| GCF_033230305.1 | HPA0170 |
| GCF_036232745.1 | F048 |
| GCF_036233215.1 | F031 |
| GCF_036233455.1 | F019 |
| GCF_036233755.1 | F004 |
| GCF_039512425.1 | PAE3 |
| GCF_039908075.1 | FIDG-26323 |
| GCF_042187215.1 | CW13 |
| GCF_044759015.1 | PA6536 |
| GCF_047801635.1 | ST175LTH |
| GCF_049532605.1 | Paer10 |
| GCF_049912995.1 | Paer99 |
| GCF_049913045.1 | Paer112 |
| GCF_050238075.1 | A27087 |
| GCF_000014625.1 | UCBPP-PA14 |
| GCF_000168335.1 | PACS2 |
| GCF_000359505.1 | B136-33 |
| GCF_000481885.1 | CF5 |
| GCF_000504045.1 | MTB-1 |

**Table 4.** NCBI accession numbers and corresponding *Mycobacterium tuberculosis* strain names.

| NCBI accession | strain name |
| --- | --- |
| GCF_000195955.2 | **H37Rv** |
| GCF_000234725.1 | Mexico |
| GCF_000572125.1 | HKBS1 |
| GCF_000934325.3 | SP38 |
| GCF_001708265.1 | SCAID 252.0 |
| GCF_001938725.1 | H37Ra |
| GCF_002447075.1 | MDRDM260 |
| GCF_002447275.1 | SLM040 |
| GCF_002447695.1 | LN3588 |
| GCF_002447715.1 | LN3589 |
| GCF_002447735.1 | LN3668 |
| GCF_002447875.1 | MDRMA203 |
| GCF_002448095.1 | TBDM2487 |
| GCF_002886165.1 | GG-5-10 |
| GCF_002886225.1 | GG-77-11 |
| GCF_002886585.1 | GG-229-10 |
| GCF_002887255.1 | GG-109-10 |
| GCF_002975475.1 | 2002/0476 |
| GCF_003265005.1 | RUS_B0 |
| GCF_009762675.2 | TCDC11 |
| GCF_013010385.1 | 4860 |
| GCF_014899525.1 | SEA11020092P6C4 |
| GCF_014899765.1 | SEA00042P6C4 |
| GCF_014899825.1 | 4-0077P6C4 |
| GCF_014900295.1 | 2-0021P6C4 |
| GCF_014900335.1 | 1-0168P6C4 |
| GCF_014900355.1 | 1-0160P6C4 |
| GCF_014900555.1 | 1-0107P6C4 |
| GCF_014900795.1 | 1-0044P6C4 |
| GCF_016917775.1 | 11502 |
| GCF_022871125.1 | SGF0472017 |
| GCF_028583185.1 | Mb3114 |
| GCF_029867205.1 | I0003758-5 |
| GCF_030574935.1 | MTb-Oman-3212387 |
| GCF_037101195.1 | DN181 |
| GCF_039907365.1 | 9 |
| GCF_044359985.1 | AST-T2 |
| GCF_051294285.1 | LP-0474076-RM2 |
| GCF_051294335.1 | LP-0474076-RM2 |
| GCF_051294345.1 | LP-0261867-RM5 |
| GCF_963993175.1 | P007-N1108 |
| GCF_963993195.1 | P008-N1380 |
| GCF_963993255.1 | P059-N1392 |
| GCF_965122915.1 | isolate WBB1009_SL1875 |
| GCF_965124535.1 | isolate WBB447_G67 |
| GCF_977011255.1 | K02C1229 |
| GCF_977011275.1 | K48KI078 |
| GCF_977011425.1 | K41T0787 |
| GCF_977011495.1 | K17T0895 |
| GCF_977011525.1 | K56TAT130 |

## B Threshold Selection

We describe how the thresholds *t*_1_ (for Jaccard similarity) and *t*_2_ (for the normalized alignment score) were selected for the annotation harmonization preprocessing step. Our goal is twofold: first, to illustrate how the same thresholds behave across different bacterial species, and second, to demonstrate that these thresholds, while not necessarily optimal for every organism, can still perform reasonably well in practice. To evaluate the thresholds, we analyzed datasets consisting of 50 strains of *Pseudomonas aeruginosa* (Table 3) and 50 strains of *Mycobacterium tuberculosis* (Table 4).

The first threshold *t*_1_ is applied to the Jaccard similarity between sets of *k*-mers extracted from protein sequences. Importantly, no value greater than zero can guarantee capturing all protein pairs that belong to the same gene group. Proteins annotated with the same gene name may still have a Jaccard similarity of zero if they share no *k*-mers, even for relatively small *k* values such as *k* = 6 or *k* = 5. However, this issue is addressed during the annotation harmonization, where proteins with the same gene name are grouped together even if they do not pass the first or second threshold tests.

If one protein sequence is a (short) substring of another, the Jaccard similarity can still be low when there is a large difference in sequence length, even though all *k*-mers of the shorter sequence are contained in the longer one. Consequently, *t*_1_ must be chosen low enough to retain biologically related proteins while avoiding an excessive number of pairwise alignments in the subsequent step.

To guide this choice, we examined the distribution of Jaccard similarities for all protein pairs in *P. aeruginosa* and *M. tuberculosis* (Figure 6). These histograms show that the vast majority of protein pairs have a similarity close to zero. For example, in the *Pseudomonas* dataset (Figure 6), more than 99.97% of values fall into the first bin (0–0.02). Therefore, choosing *t*_1_ = 0.02 would be a good choice for limiting the number of alignments required in the next step. We later show that this threshold is also adequate for retaining most biologically relevant protein pairs.

**Figure 6.**
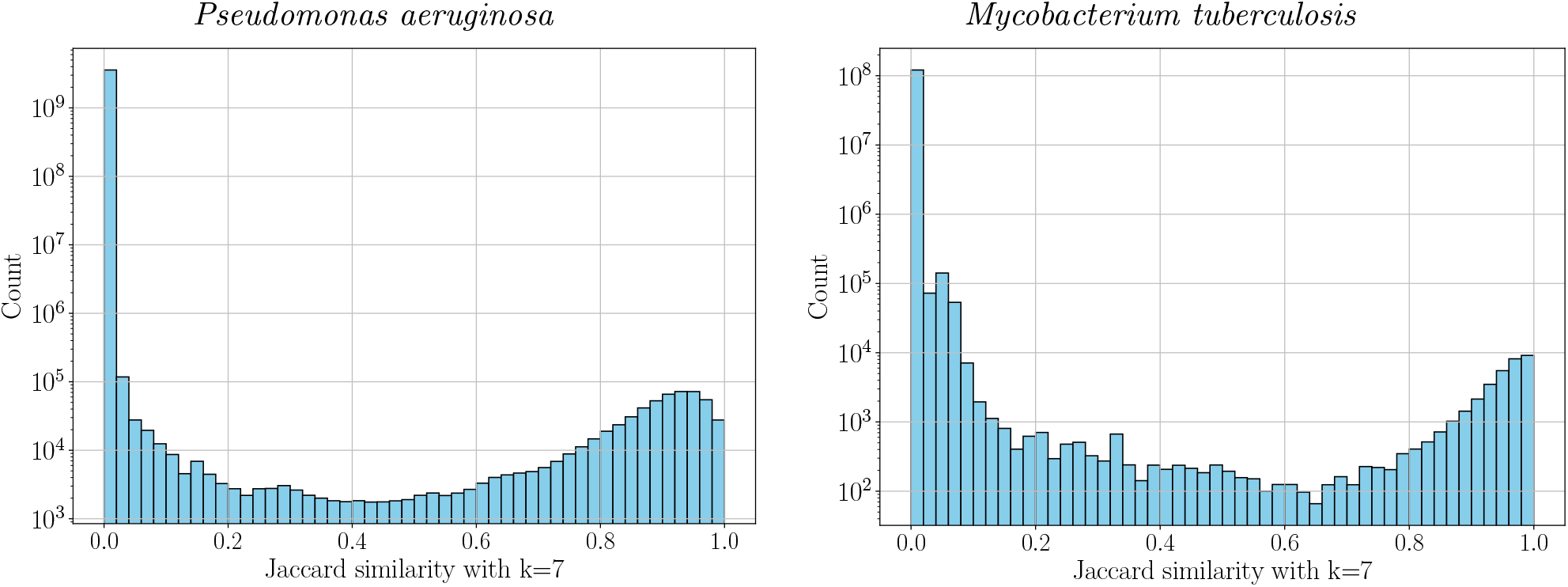
Histogram of Jaccard similarities between sets of amino acid 7-mers of all protein pairs across 50 strains of *Pseudomonas aeruginosa* (left) and *Mycobacterium tuberculosis*. Self-similarities (i.e., comparisons of a protein with itself) are excluded.

The second threshold *t*_2_ is applied to the normalized alignment score obtained from pairwise sequence alignments. High values of *t*_2_ ensure that grouped proteins correspond to the same gene but may exclude proteins that share highly similar regions with the group, potentially resulting in ambiguous mapping during read-to-gene mapping. Conversely, lower *t*_2_ values reduce ambiguous mapping by allowing reads to be assigned to larger gene groups, but at the cost of reduced specificity in gene classification and expression estimation. Figure 7 shows the distribution of normalized alignment scores for all protein pairs with Jaccard similarity above *t*_1_ = 0.02.

**Figure 7.**
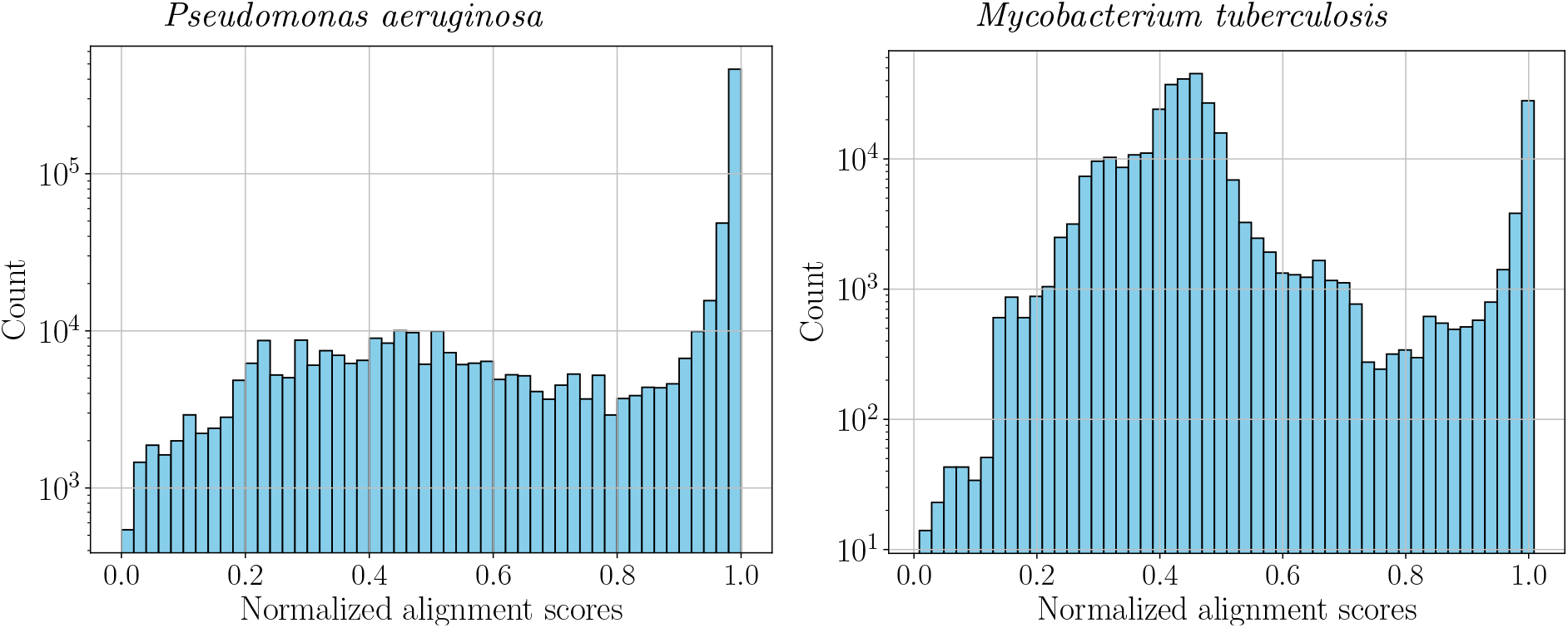
Histogram of the normalized alignment score for all protein pairs with Jaccard similarities between sets of amino acid 7-mers above *t*_1_ = 0.02 across 50 strains of *Pseudomonas aeruginosa* (left) and *Mycobacterium tuberculosis*

To evaluate how *t*_1_ and *t*_2_ affect read mapping, we examined precision (fraction of correctly mapped reads among mapped reads) and fraction of ambiguously mapped reads for different threshold combinations using simulated reads. The simulation followed the procedure described in the simulated data results section of the main article, using single-end reads and a reduced average coverage of 20.

Mapping these reads with PanXpress without the correction of the annotations resulted in approximately 5% of ambiguously mapped reads, which reduced precision and worsened the gene expression estimate. Figure 8 shows the fraction of ambiguously mapped reads for different threshold combinations and two distinct species, while Figure 9 shows the false mapping rates, defined as 1 *−* precision.

**Figure 8.**
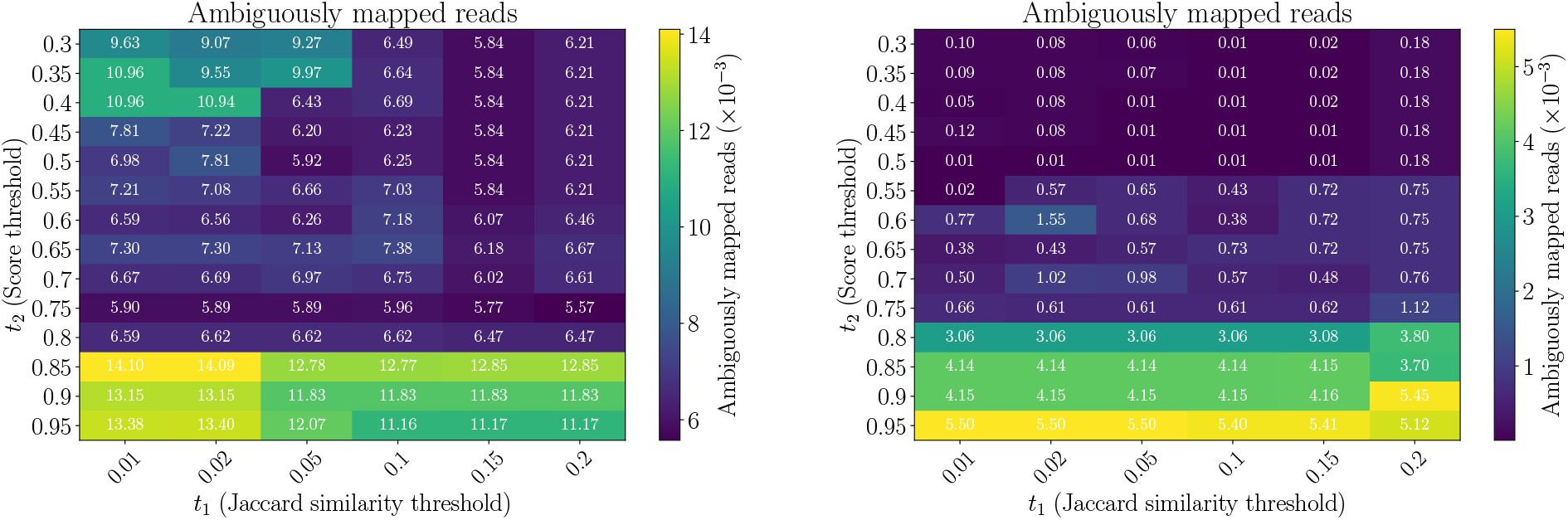
Proportion of ambiguously mapped reads using simulated reads from 3 strains of *Pseudomonas aeruginosa* (left) and *Mycobacterium tuberculosis* (right) using a reference of 50 strains of the same species (coverage 20x, read length 100, 15% upregulated genes, 15% downregulated genes).

**Figure 9.**
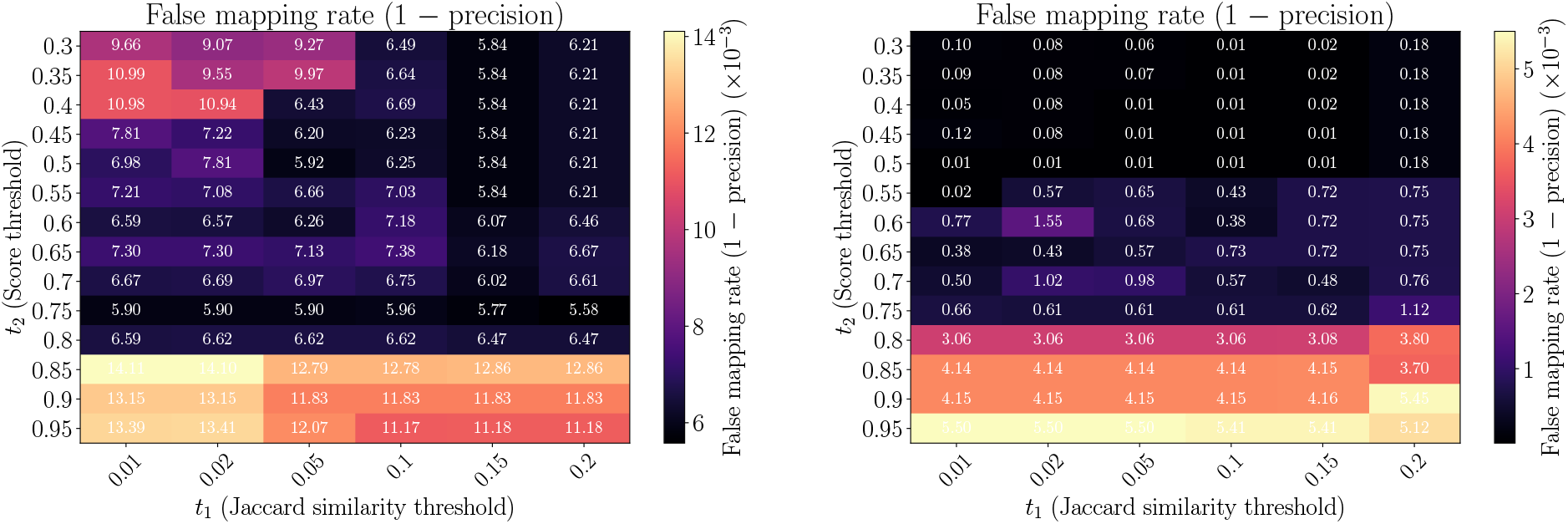
False mapping rate (1*−* precision) using simulated reads from 3 strains of *Pseudomonas aeruginosa* (left) and *Mycobacterium tuberculosis* (right) using a reference of 50 strains of the same species (coverage 20x, read length 100, 15% upregulated genes, 15% downregulated genes).

Figures 8 and 9 show that a decrease in the proportion of ambiguously mapped reads is generally correlated with a decrease in precision error. In most cases, lowering the Jaccard similarity threshold *t*_1_ reduces ambiguous mapping and improves precision, or at least does not worsen these metrics. An exception occurs for very small values of *t*_2_ in the *Pseudomonas* dataset, where decreasing *t*_1_ can slightly worsen the results. This observation suggests that extremely low values of *t*_2_ should be avoided when selecting general thresholds applicable across species.

We also observed that reducing *t*_1_ from 0.02 to 0.01 results in only marginal improvements in ambiguous mapping and precision. Given the substantial increase in the number of pairwise alignments required at lower thresholds, *t*_1_ = 0.02 represents a reasonable compromise for computational efficiency.

Based solely on the mapping metrics shown in Figures 8 and 9, one might conclude that, for a fixed *t*_1_, the score threshold *t*_2_ should be chosen as low as possible to minimize ambiguous mapping. However, low values of *t*_2_ lead to a significant loss of specificity by merging distinct genes into the same group. For example, in the *Pseudomonas* dataset, using *t*_1_ = 0.01 and *t*_2_ = 0.5 produces gene groups with names such as `fimU_gspG_gspH_pilA_pilE_xcpT_xcpU_xcpW` or `phzA_phzA1_phzA2_phzB_phzB1_phzB2`. Although these genes may share a common evolutionary origin and therefore achieve moderate alignment scores, merging them into a single group is undesirable for downstream analysis.

Increasing the threshold to *t*_2_ = 0.75 separates several of these merged families, for example, splitting the `phzA` and `phzB` variants into distinct groups. This improves the specificity of gene classification while still reducing ambiguous mapping.

Since the primary goal of the preprocessing step is to cluster nearly identical proteins that have been inconsistently annotated across strains rather than to group distantly related genes with a common evolutionary origin, we recommend *t*_2_ = 0.75 as a practical compromise between specificity and mapping performance.

At thresholds *t*_1_ = 0.02, *t*_2_ = 0.75, the proportion of ambiguously mapped reads is approximately 0.59% for *Pseudomonas aeruginosa* and 0.06% for *Mycobacterium tuberculosis*, which is nearly an order of magnitude lower than when no re-annotation is applied (for *Pseudomonas*). As shown in the simulated data results section, these thresholds are sufficient to obtain highly accurate gene expression estimates.

We acknowledge that these thresholds may not be optimal for all organisms or datasets. However, our results demonstrate that they effectively reduce ambiguously mapped reads caused by inconsistent gene annotations while assigning the same gene name to nearly identical proteins across strains. Overall, the selected thresholds provide a reasonable and robust choice across multiple bacterial species.

## C Pangenome analysis

We examine how the number of unique gene names evolves as additional strains are included. Figure 10 show the increase of unique well-defined gene names (ignoring hypothetical proteins) for increasing numbers of strains before resolving annotation inconsistencies. To generate these plots, we randomly ordered the 50 strains for each species while placing the reference strain first in the list. For each value *s* of strains (x-axis), we considered the first *s* strains and counted the number of unique gene names present. For *P. aeruginosa*, the number of unique gene names increases only slowly between 30 and 50 strains. For *M. tuberculosis*, the difference between using 10 and 50 strains is barely noticeable, suggesting that approximately 10 strains already capture most of the gene name diversity.

**Figure 10.**
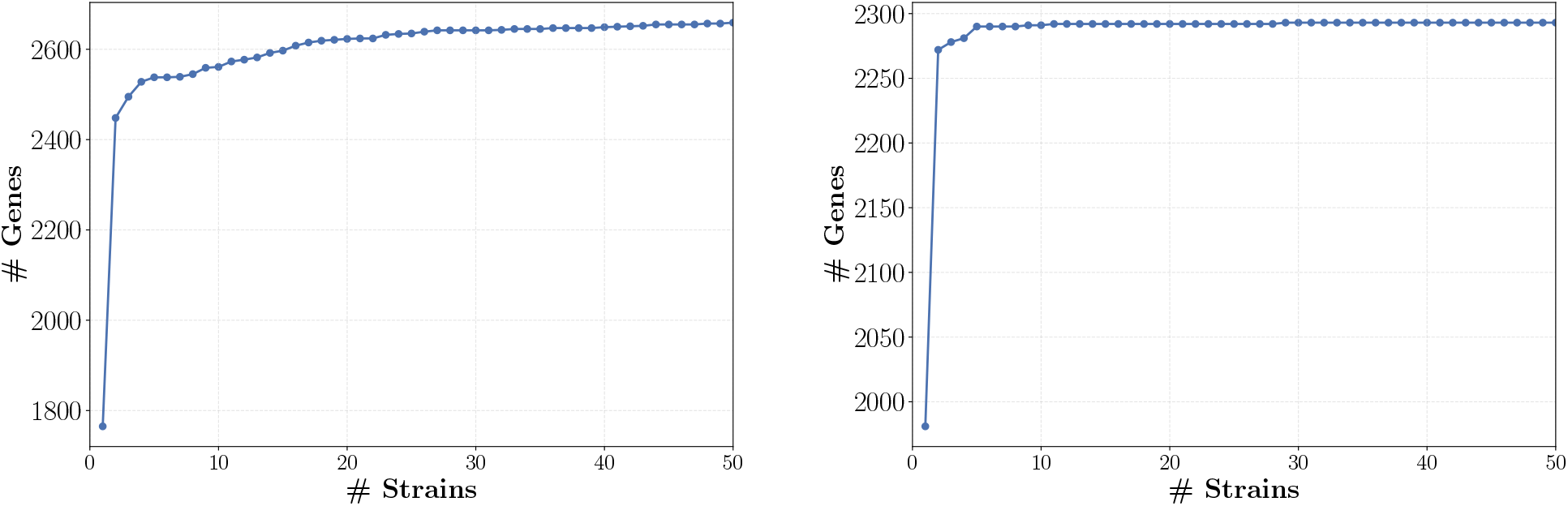
Number of unique gene names in 50 strains of *P. aeruginosa* (left) and *M*.*tuberculosis* (right) before annotation harmonization.

The reference strains typically have highly curated annotation files. As a result, some coding regions annotated with well-defined gene names in the reference strain may be labeled as hypothetical proteins in other strains and are therefore ignored in this analysis. Additionally, distinct paralogs may have different names in the reference strain. These factors may explain why adding more strains introduces relatively little new information in terms of additional gene names after the first few strains are included.

We examined the distribution of the number of colors (genes) associated with indexed *k*-mers for *P. aeruginosa*. Here we perform the same analysis for *M. tuberculosis*, shown in Figure 11.

**Figure 11.**
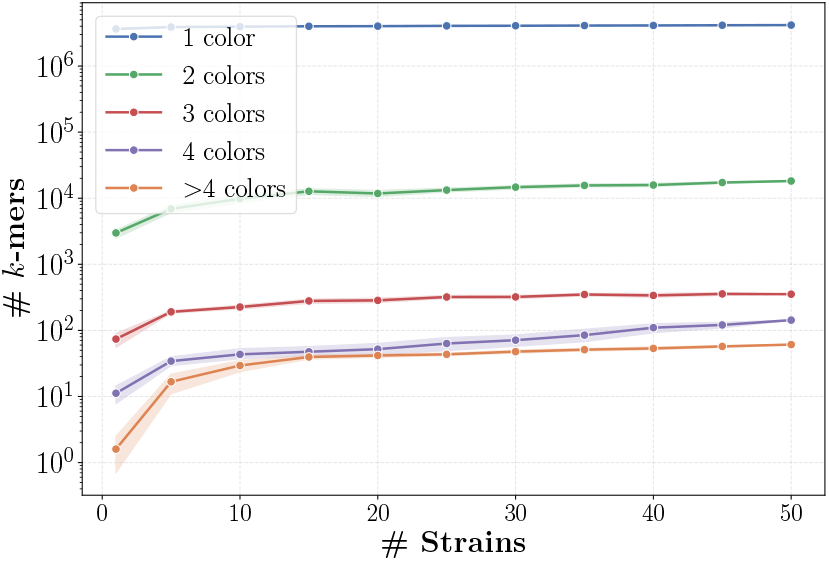
Number of *k*-mers associated with different numbers of colors in the PanXpress index as a function of the number of strains of *M. tuberculosis*. For each number of strains, the experiment was repeated 10 times using randomly sampled strains. Lines represent the mean number of *k*-mers and the shaded areas indicate 95% confidence intervals.

As observed for *P. aeruginosa*, most *k*-mers are unique to a single gene. However, in the *M. tuberculosis* dataset, the number of *k*-mers associated with more than four colors is larger than in the *P. aeruginosa* dataset. Nevertheless, this number remains below 100, which is negligible compared to the total number of indexed *k*-mers (in the millions).

We also observe that after including approximately 10 strains, only a small number of additional *k*-mers are introduced with each new strain. This behavior is consistent with the observed saturation in the number of unique gene names.

## D Fold change analysis for simulated data

We present further results regarding the comparison between the experimental fold changes obtained with each tool and the expected fold changes for the simulated reads described in the Results section.

Figure 12 compares the experimental fold changes obtained with each tool to the expected fold changes for simulated single-end reads from *Pseudomonas aeruginosa*. The close agreement between the experimental and expected fold changes indicates that all four tools accurately capture the differential expression trends.

**Figure 12.**
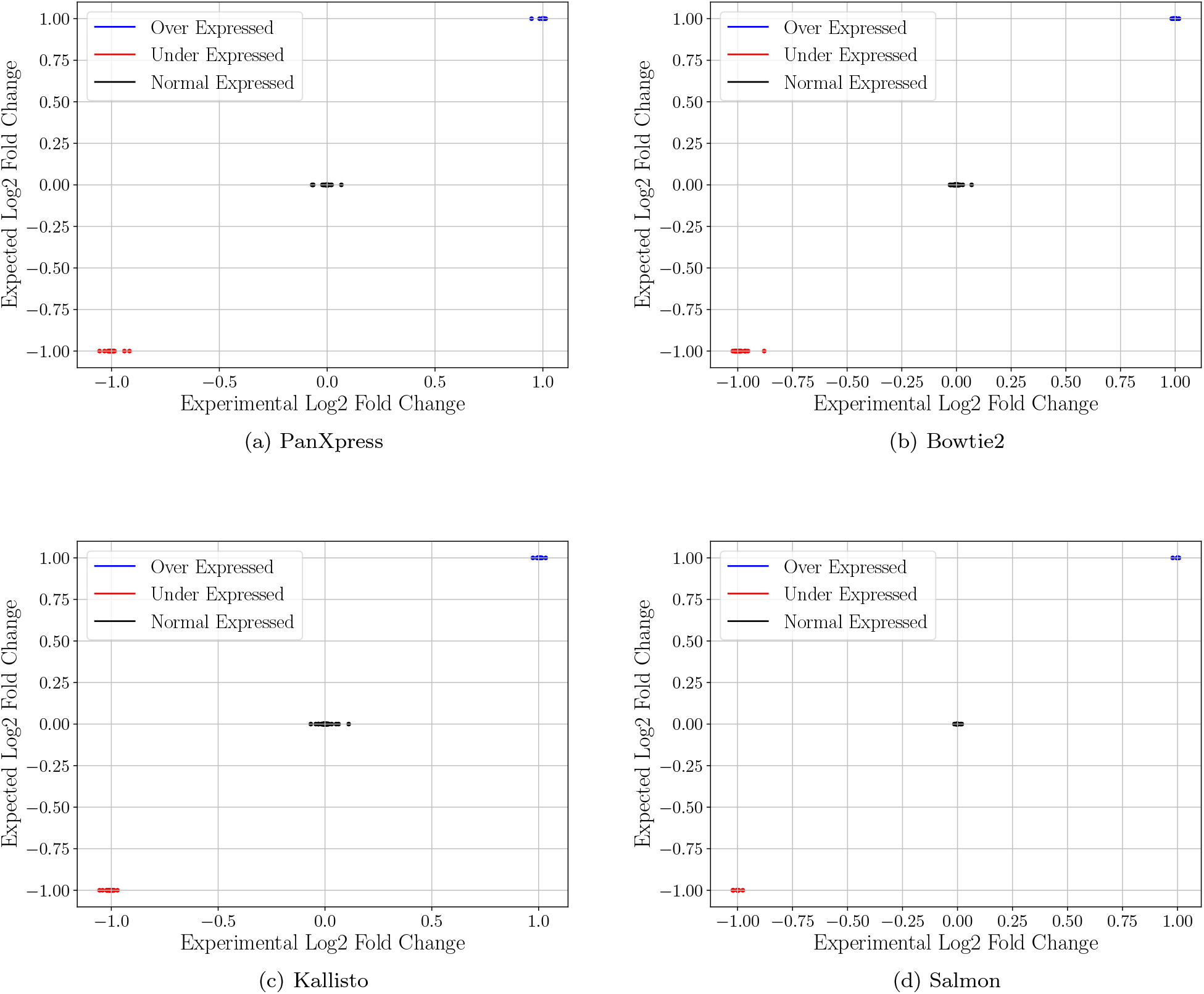
Comparison between expected and experimentally estimated gene expression fold changes for simulated single-end reads. Expected true log_2_ fold changes were at *−*1 (15% of genes), +1 (15% of genes) and 0 (remaining 70% of genes).

Since the results for paired-end data are very similar, the corresponding figures are omitted for brevity. The same results are observed for *Mycobacterium tuberculosis*.

We performed the same comparative analysis for *Mycobacterium tuberculosis* as previously done for Pseudomonas aeruginosa. Specifically, we evaluated read mapping performance of PanXpress and Bowtie2 in terms of precision and sensitivity (Table 5). Results show the same overall conclusions as observed for *Pseudomonas aeruginosa*. PanXpress achieves very high precision and recall, with recall being slightly lower than precision. This is expected because PanXpress performs confident mapping: Reads are only assigned when the mapping is unambiguous. As a consequence, PanXpress achieves higher precision than Bowtie2, while a small fraction of reads remain unmapped, resulting in a slightly lower recall.

**Table 5.** Read mapping performance comparison between PanXpress and Bowtie2 on simulated *M. tuberculosis* reads, in single-end (SE) and paired-end (PE) mode. Results are reported for two independent control samples (control 1, 2) and two independent samples with modified expression (up-down 1, 2). Precision (Prec%): correctly mapped reads of mapped reads. Recall (Recall%): correctly mapped reads of all reads.

| Dataset | PanXpress |  | Bowtie2 |  |
| --- | --- | --- | --- | --- |
|  | Prec% | Recall% | Prec% | Recall% |
| SE control 1 | 100.000 | 99.922 | 99.919 | 99.919 |
| SE control 2 | 100.000 | 99.922 | 99.920 | 99.920 |
| SE up-down 1 | 100.000 | 99.929 | 99.921 | 99.921 |
| SE up-down 2 | 100.000 | 99.930 | 99.922 | 99.922 |
| PE control 1 | 100.000 | 99.943 | 99.944 | 99.944 |
| PE control 2 | 100.000 | 99.941 | 99.946 | 99.946 |
| PE up-down 1 | 100.000 | 99.946 | 99.948 | 99.948 |
| PE up-down 2 | 100.000 | 99.947 | 99.947 | 99.947 |

Furthermore, we assessed quantification accuracy across PanXpress, Salmon, Kallisto, and Bowtie2 using root mean squared error (RMSE) and mean absolute error (MAE) of estimated versus true gene expression fold changes (Table 6). These analyses were conducted for both single-end (SE) and paired-end (PE) simulated reads. Again, we see results similar to those observed for *Pseudomonas aeruginosa*. Salmon achieves the lowest errors for single-end reads, while PanXpress yields the lowest errors for paired-end reads. In general, PanXpress and Salmon produce substantially lower quantification errors than Bowtie2. Additionally, in contrast to the results obtained for P. *aeruginosa*, Kallisto achieves a smaller RMSE than Bowtie2 in this dataset.

**Table 6.**
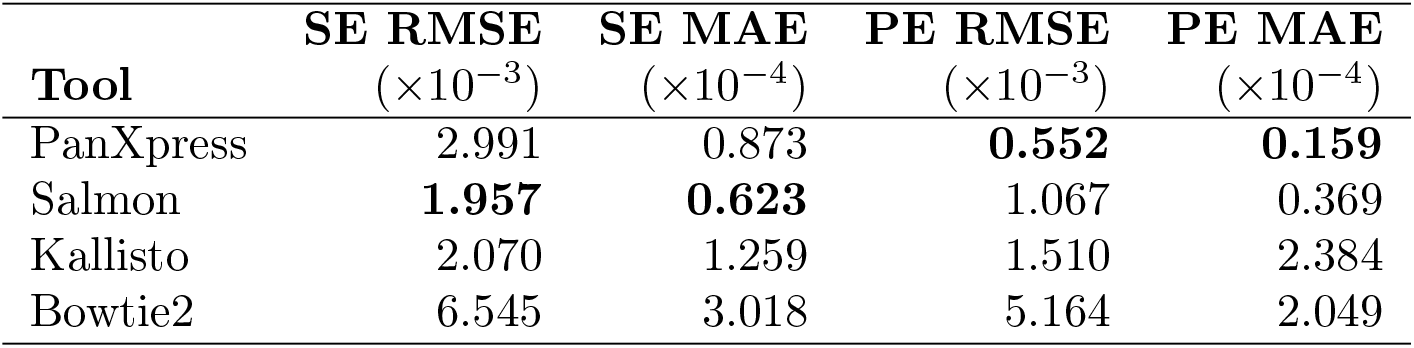
Quantification error comparison for PanXpress, Salmon, Kallisto, and Bowtie2 on simulated *M. tuberculosis* single-end (SE) and paired-end (PE) reads, measured as root mean squared error (RMSE) and mean absolute error (MAE) between estimated and true gene expression log_2_ fold changes.

| Tool | SE RMSE | SE MAE | PE RMSE | PE MAE |
| --- | --- | --- | --- | --- |
| | ($\times 10^{-3}$) | ($\times 10^{-4}$) | ($\times 10^{-3}$) | ($\times 10^{-4}$) |
| PanXpress | 2.991 | 0.873 | **0.552** | **0.159** |
| Salmon | **1.957** | **0.623** | 1.067 | 0.369 |
| Kallisto | 2.070 | 1.259 | 1.510 | 2.384 |
| Bowtie2 | 6.545 | 3.018 | 5.164 | 2.049 |

